## Supplementary File 1 for "An adjunctive therapy administered with an antibiotic prevents enrichment of antibiotic-resistant clones of a colonizing opportunistic pathogen"

**Model 1: Effect of daptomycin on proportion of daptomycin-resistant VR *E. faecium***

**Input data:** Fig2AB_VRESusceptibility.csv on Dryad

**Model structure in R:**
glmmTMB(CFU_DAP ~ Antibiotic + Day + offset(log(CFU_noDAP)) + (1|Mouse), data = na.omit(data), ziformula = ~1, family = nbinom1)

summary()

| | Estimate | Std. Error | z value | Pr(>\|z\|) |
|---|---|---|---|---|
| (Intercept) | -4.581731 | 0.365351 | -12.541 | < 2e-16 |
| Antibioticdaptomycin | 4.645379 | 0.365523 | 12.709 | < 2e-16 |
| Day | -0.032638 | 0.008015 | -4.072 | 4.66e-05 |

**Model 2: Effect of daptomycin on total shedding of VR *E. faecium***

**Input data:** SuppFig1_VREDensity.csv on Dryad

**Model structure in R:**
glmmTMB(VRE.CFU.per.10mgFeces ~ Antibiotic + Day + (1|Mouse), data = data[data$Day>1,], ziformula=~1, family=nbinom1)

| | Estimate | Std. Error | z value | Pr(>\|z\|) |
|---|---|---|---|---|
| (Intercept) | 13.117624 | 0.441897 | 29.685 | <2e-16 |
| Antibioticdaptomycin | 0.001027 | 0.403890 | 0.003 | 0.9980 |
| Day | 0.045302 | 0.022206 | 2.040 | 0.0413 |

**Model 3: Effect of cholestyramine on proportion of daptomycin-resistant VR *E. faecium***

**Input data:** Fig5_VREDensity.csv on Dryad

**Data processing:**
1. Create a version of Day variable structured as a factor with levels 1:14 (Day.factor).
2. Drop entries where noDAP.VRE.CFU.per.10mgFeces < 20 (detection limit).
3. Create a variable for proportion resistant colonies (DAP.VRE.CFU.per.10mgFeces /noDAP.VRE.CFU.per.10mgFeces)
4. Normalize proportions to be bounded by 0 and 1. Divide all proportions by maximum proportion value, to create variable PropNorm.

**Model structure in R:**

glmmTMB(PropNorm ~ Diet*Antibiotic + Day + Experiment + (1|Mouse) + ar1(Day.factor + 0 | Mouse), data = data[data$Day>1,]), family = binomial, weights=

noDAP.VRE.CFU.per.10mgFeces, control = glmmTMBControl(optimizer = optim, optArgs = list(method="BFGS")))

**Output:**
summary()

| | Estimate | Std. Error | z value | Pr(>\|z\|) |
|---|---|---|---|---|
| (Intercept) | -6.21957 | 0.50384 | -12.344 | < 2e-16 |
| Dietcholestyramine | 0.31196 | 0.55510 | 0.562 | 0.57413 |
| Antibioticdaptomycin | 4.24574 | 0.46674 | 9.097 | < 2e-16 |
| Day | 0.01062 | 0.02381 | 0.446 | 0.65556 |
| ExperimentB | 0.66805 | 0.44487 | 1.502 | 0.13319 |
| ExperimentC | -3.53285 | 0.40434 | -8.737 | < 2e-16 |
| ExperimentD | -1.79728 | 0.45953 | -3.911 | 9.19e-05 |
| Dietcholestyramine: Antibioticdaptomycin | -2.16041 | 0.65702 | -3.288 | 0.00101 |

Random effects:

| Groups | Name | Variance | Std.Dev. | Corr |
|---|---|---|---|---|
| Mouse | (Intercept) | 4.130e-07 | 0.0006427 | |
| Mouse.1 | Day.factor2 | 6.434e+00 | 2.5366239 | 0.54 (ar1) |

Note: We also ran this model including only data with a higher total number of bacterial colonies counted (noDAP.VRE.CFU.per.10mgFeces > 150). This was a check to see if the results were affected by samples with low total counts, which might give inaccurate proportions. The key result from this model, a significant effect of Diet*Antibiotic, remains true with the higher cutoff (p = 0.0004).

**Model 4: Effect of cholestyramine on proportion of daptomycin-resistant VR *E. faecium*, Days 8 & 14**

**Input data:** Fig5_VRESusceptibility.csv on Dryad

**Data processing:**
1. Create a variable for proportion resistant colonies (CFU_DAP/CFU_noDAP).
3. Normalize proportions to be bounded by 0 and 1. Divide all proportions by maximum proportion value, to create variable PropNorm.

**Model structure in R:**

glmmTMB(PropNorm ~ Diet*Antibiotic + Day + Experiment + (1|Mouse), data = na.omit(data), family = binomial, weights = CFU_noDAP)

summary()

| | Estimate | Std. Error | z value | Pr(>\|z\|) |
|---|---|---|---|---|
| (Intercept) | -7.20705 | 0.70739 | -10.188 | < 2e-16 |
| Dietcholestyramine | 1.22512 | 0.84030 | 1.458 | 0.145 |
| Antibioticdaptomycin | 6.46804 | 0.72579 | 8.912 | < 2e-16 |
| Day14 | 0.12750 | 0.02077 | 6.140 | 8.27e-10 |

| | | | | |
|---|---|---|---|---|
| ExperimentB | 0.78377 | 0.57735 | 1.358 | 0.175 |
| ExperimentC | -4.12226 | 0.60214 | -6.846 | 7.59e-12 |
| ExperimentD | -0.13626 | 0.61947 | -0.220 | 0.826 |
| Dietcholestyramine: Antibioticdaptomycin | -4.06349 | 0.97461 | -4.169 | 3.05e-05 |

Random effects:

| Groups | Name | Variance | Std.Dev. |
|---|---|---|---|
| Mouse | (Intercept) | 4.7 | 2.168 |

**Model 5: Effect of cholestyramine on shedding of daptomycin-resistant VR *E. faecium***

**Input data:** Fig5_VREDensity.csv on Dryad

**Data processing:** Added 1 to density variables DAP.VRE.CFU.per.10mgFeces and noDAP.VRE.CFU.per.10mgFeces so that data could be logged.

**Model structure in R:**

```
lme(data=data[data$Day >1,],
    log10(DAP.VRE.CFU.per.10mgFeces) ~ Antibiotic*Diet* Day + Experiment,
    random=~1|Mouse, correlation = corAR1(form = ~Day|Mouse))
```

**Output:**

anova()

| | **numDF** | **denDF** | **F-value** | **p-value** |
|---|---|---|---|---|
| (Intercept) | 1 | 556 | 1189.4014 | <.0001 |
| Antibiotic | 1 | 133 | 3.4018 | 0.0674 |
| Diet | 1 | 133 | 1.0057 | 0.3178 |
| Day | 1 | 556 | 27.6576 | <.0001 |
| Experiment | 3 | 133 | 32.1013 | <.0001 |
| Antibiotic:Diet | 1 | 133 | 0.0993 | 0.7532 |
| Antibiotic:Day | 1 | 556 | 25.4397 | <.0001 |
| Diet:Day | 1 | 556 | 8.1842 | 0.0044 |
| Antibiotic:Diet:Day | 1 | 556 | 23.2725 | <.0001 |

summary()

| | **Value** | **Std. Error** | **DF** | **t-value** | **p-value** |
|---|---|---|---|---|---|
| (Intercept) | 5.697248 | 0.4507439 | 556 | 12.639655 | 0.0000 |
| Antibioticdaptomycin | -1.241995 | 0.4711206 | 133 | -2.636257 | 0.0094 |
| Dietcholestyramine | -0.989360 | 0.5613877 | 133 | -1.762348 | 0.0803 |
| Day | -0.249036 | 0.0370298 | 556 | -6.725290 | 0.0000 |
| ExperimentB | 1.019619 | 0.3543514 | 133 | 2.877423 | 0.0047 |
| ExperimentC | -1.884019 | 0.3188225 | 133 | -5.909305 | 0.0000 |
| ExperimentD | -1.058643 | 0.3543514 | 133 | -2.987552 | 0.0033 |
| Antibioticdaptomycin:Dietcholestyramine | 1.841121 | 0.6642429 | 133 | 2.771759 | 0.0064 |
| Antibioticdaptomycin:Day | 0.304636 | 0.0438142 | 556 | 6.952911 | 0.0000 |
| Dietcholestyramine:Day | 0.129620 | 0.0523680 | 556 | 2.475173 | 0.0136 |

| | | | | | |
|---|---|---|---|---|---|
| Antibioticdaptomycin:Dietcholestyramine:Day | -0.293358 | 0.0619626 | 556 | -4.734433 | 0.0000 |

Random effects:
Formula: ~1 | Mouse

| | (Intercept) | Residual |
|---|---|---|
| StdDev: | 1.19095 | 1.524979 |

**Model 6: Effect of cholestyramine on total VR *E. faecium* shedding**

**Input data:** Fig5_VREDensity.csv on Dryad

**Data processing:** Added 1 to density variables DAP.VRE.CFU.per.10mgFeces and noDAP.VRE.CFU.per.10mgFeces so that data could be logged.

**Model structure in R:**

```
lme(data=data[data$Day >1,],
    log10(noDAP.VRE.CFU.per.10mgFeces) ~ Antibiotic*Diet* Day + Experiment,
    random=~1|Mouse, correlation = corAR1(form = ~Day|Mouse))
```

**Output:**

anova()

| | numDF | denDF | F-value | p-value |
|---|---|---|---|---|
| (Intercept) | 1 | 556 | 1893.8435 | <.0001 |
| Antibiotic | 1 | 133 | 6.5129 | 0.0118 |
| Diet | 1 | 133 | 1.1657 | 0.2822 |
| Day | 1 | 556 | 26.5513 | <.0001 |
| Experiment | 3 | 133 | 5.9143 | 0.0008 |
| Antibiotic:Diet | 1 | 133 | 1.5648 | 0.2132 |
| Antibiotic:Day | 1 | 556 | 2.7475 | 0.0980 |
| Diet:Day | 1 | 556 | 16.0355 | 0.0001 |
| Antibiotic:Diet:Day | 1 | 556 | 14.3794 | 0.0002 |

summary()

| | Value | Std. Error | DF | t-value | p-value |
|---|---|---|---|---|---|
| (Intercept) | 7.144392 | 0.4763063 | 556 | 14.999573 | 0.0000 |
| Antibioticdaptomycin | -2.095335 | 0.4937631 | 133 | -4.243604 | 0.0000 |
| Dietcholestyramine | -0.600540 | 0.5882331 | 133 | -1.020921 | 0.3091 |
| Day | -0.134885 | 0.0365464 | 556 | -3.690774 | 0.0002 |
| ExperimentB | 0.722889 | 0.3851389 | 133 | 1.876956 | 0.0627 |
| ExperimentC | -0.614066 | 0.3465231 | 133 | -1.772078 | 0.0787 |
| ExperimentD | -0.585066 | 0.3851389 | 133 | -1.519103 | 0.1311 |
| Antibioticdaptomycin:Dietcholestyramine | 2.275048 | 0.6960068 | 133 | 3.268715 | 0.0014 |
| Antibioticdaptomycin:Day | 0.166631 | 0.0432423 | 556 | 3.853423 | 0.0001 |
| Dietcholestyramine:Day | 0.055012 | 0.0516845 | 556 | 1.064374 | 0.2876 |

| | | | | | |
|---|---|---|---|---|---|
| Antibioticdaptomycin:Dietcholestyramine :Day | -0.231896 | 0.0611539 | 556 | -3.792015 | 0.0002 |

Random effects:
Formula: ~1 | Mouse

| | (Intercept) | Residual |
|---|---|---|
| StdDev: | 1.331139 | 1.505074 |

**<u>Model 7. Effect of cholestyramine on total VR *E. faecium* shedding in control mice (no daptomycin)</u>**

**Input data:** Fig5_VREDensity.csv on Dryad

**Data processing:** Added 1 to density variables DAP.VRE.CFU.per.10mgFeces and noDAP.VRE.CFU.per.10mgFeces so that data could be logged.

**Model structure in R:**

```
lme(data= data[data$Day>1 & data$Antibiotic=="control",],
    log10(noDAP.VRE.CFU.per.10mgFeces) ~ Diet*Day + Experiment, random=~1|Mouse,
    correlation = corAR1(form = ~Day|Mouse))
```

**Output:**

anova()

| | **numDF** | **denDF** | **F-value** | **p-value** |
|---|---|---|---|---|
| (Intercept) | 1 | 158 | 668.8815 | <.0001 |
| Diet | 1 | 35 | 0.2386 | 0.6282 |
| Day | 1 | 158 | 28.0471 | <.0001 |
| Experiment | 3 | 35 | 1.5892 | 0.2094 |
| Diet:Day | 1 | 158 | 1.8403 | 0.1768 |

summary()

| | **Value** | **Std. Error** | **DF** | **t-value** | **p-value** |
|---|---|---|---|---|---|
| (Intercept) | 7.120770 | 0.5537743 | 158 | 12.858614 | 0.0000 |
| Dietcholestyramine | -0.600540 | 0.5394026 | 35 | -1.113342 | 0.2731 |
| Day | -0.134885 | 0.0286741 | 158 | -4.704063 | 0.0000 |
| ExperimentB | 0.615201 | 0.6556186 | 35 | 0.938353 | 0.3545 |
| ExperimentC | -0.803163 | 0.6556186 | 35 | -1.225047 | 0.2287 |
| ExperimentD | -0.193794 | 0.6556186 | 35 | -0.295590 | 0.7693 |
| Dietcholestyramine:Day | 0.055012 | 0.0405513 | 158 | 1.356595 | 0.1768 |

Random effects:
Formula: ~1 | Mouse

| | (Intercept) | Residual |
|---|---|---|
| StdDev: | 1.367585 | 1.180871 |
