## Supplementary material for "An adjunctive therapy administered with an antibiotic prevents enrichment of antibiotic-resistant clones of a colonizing opportunistic pathogen": Figure Supplements

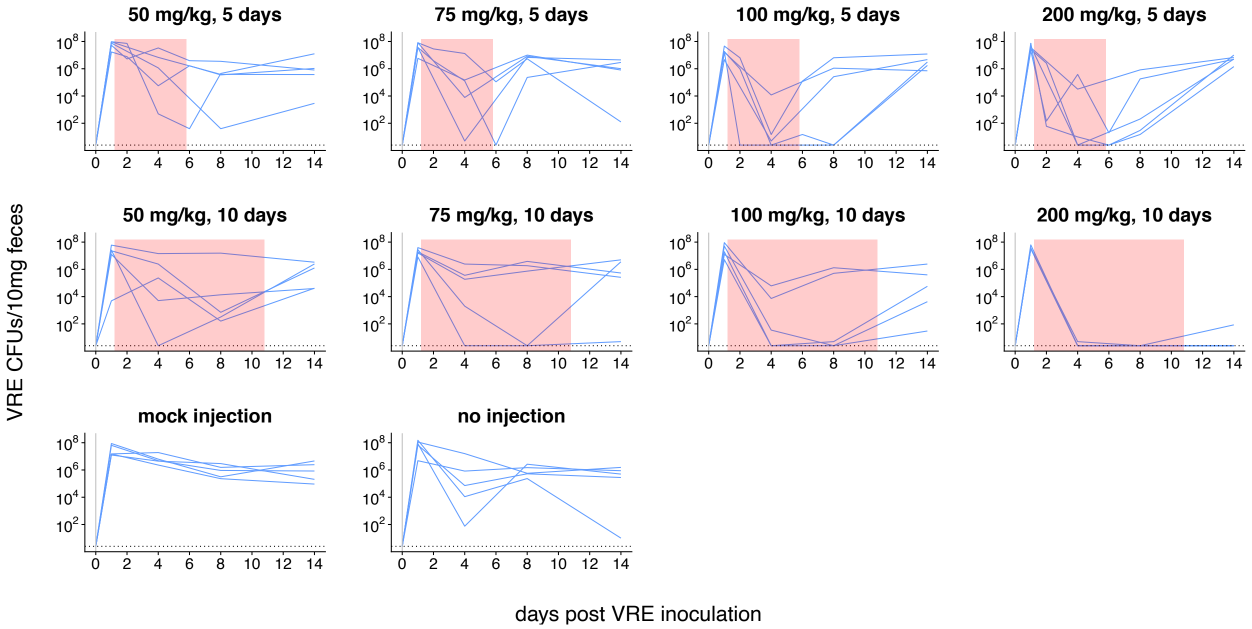


**Figure 2 - figure supplement 1.** Dynamics of VRE shedding for experiment shown in Fig. 2. Subcutaneous daptomycin dose and treatment durations are listed above each panel. Each line represents VRE densities from an individual mouse (N=5 per treatment). The pink shaded region indicates days of daptomycin therapy. The horizontal line marks the detection limit.

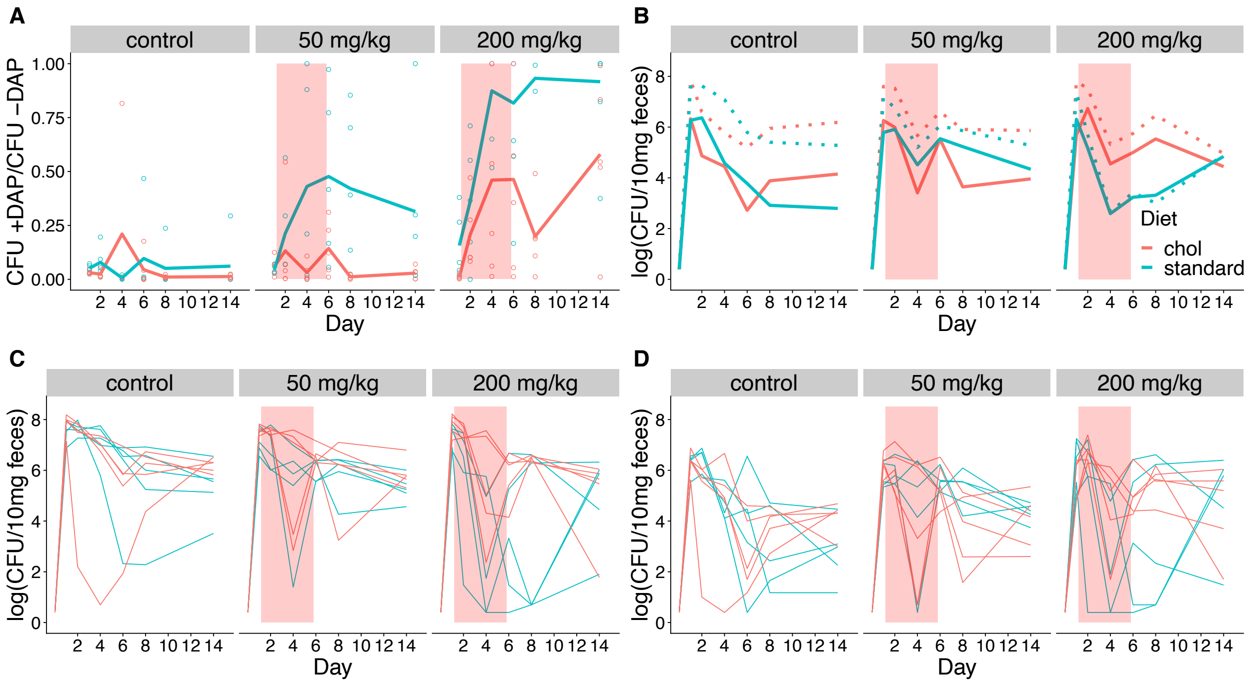


**Figure 5 - figure supplement 1.** Fecal shedding of VR E. faecium for Experiment A (strain BL00239-1, Swiss Webster mice, cholestyramine started one day prior to first daptomycin dose). (A) The proportion of fecal VR E. faecium that were daptomycin-resistant over time in mice. Proportions were determined by plating on agar with daptomycin (+DAP) and without daptomycin (-DAP). Lines show means, and open points show values for individual mice (N=5). Proportions were not determined for samples with < 20 CFU VR E. faecium per 10 mg feces, as these densities were below the limit of detection for this plating assay, and these samples were not included in Panel A. The pink shaded region indicates days of daptomycin therapy. (B) Total VR E. faecium densities corresponding to data shown in Panel A (N=5, mean shown). Dotted line shows total density (-DAP) and solid line shows the density of daptomycin-resistant VR E. faecium (+DAP). All samples, including those with low densities, were included in Panel B. (C) Total VR E. faecium densities (-DAP) for individual mice in this experiment. Each line tracks values for one mouse. (D) Daptomycin-resistant VR E. faecium densities (+DAP) for individual mice in this experiment. Each line tracks values for one mouse.

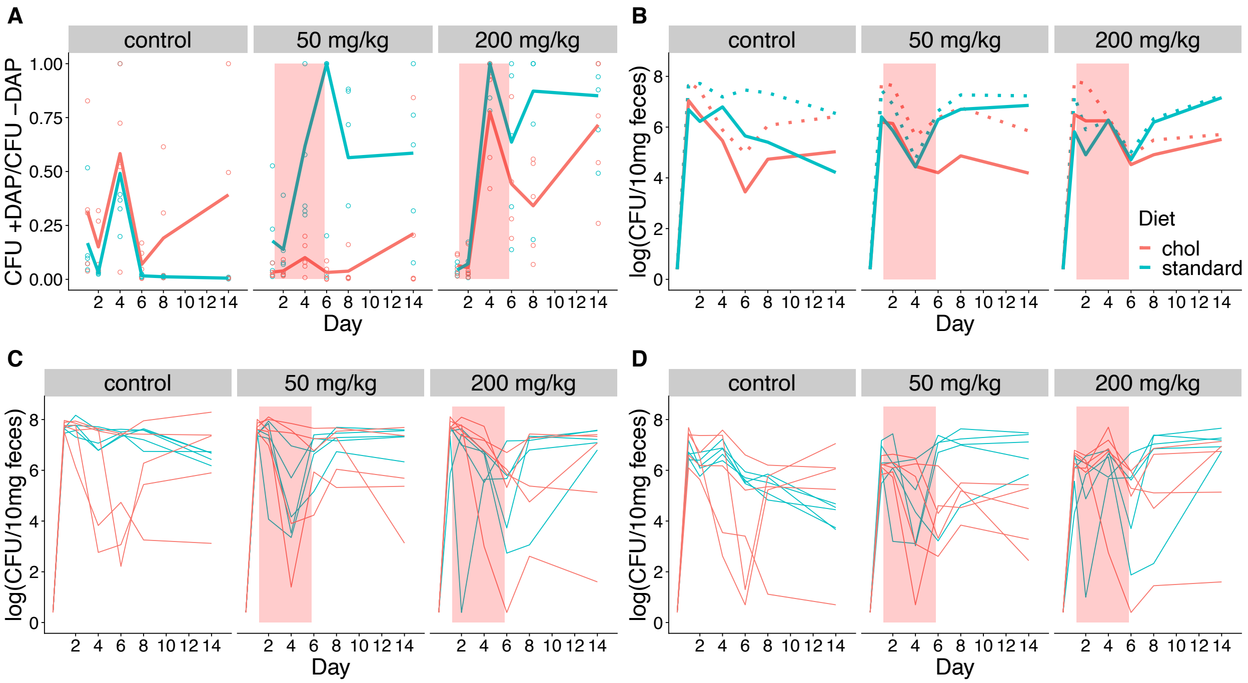


**Figure 5 - figure supplement 2.** Fecal shedding of VR *E.* faecium for Experiment B (strain BL00239-1, C57BL/6 mice, cholestyramine started one day prior to first daptomycin dose). (A) The proportion of fecal VR *E. faecium* that were daptomycin-resistant over time in mice. Proportions were determined by plating on agar with daptomycin (+DAP) and without daptomycin (-DAP). Lines show means, and open points show values for individual mice (N=5). Proportions were not determined for samples with < 20 CFU VR *E. faecium* per 10 mg feces, as these densities were below the limit of detection for this plating assay,and these samples were not included in Panel A. The pink shaded region indicates days of daptomycin therapy. (B) Total VR *E. faecium* densities corresponding to data shown in Panel A (N=5, mean shown). Dotted line shows total density (-DAP) and solid line shows the density of daptomycin-resistant VR *E. faecium* (+DAP). All samples, including those with low densities, were included in Panel B. (C) Total VR *E. faecium* densities (-DAP) for individual mice in this experiment. Each line tracks values for one mouse. (D) Daptomycin-resistant VR *E. faecium* densities (+DAP) for individual mice in this experiment. Each line tracks values for one mouse.

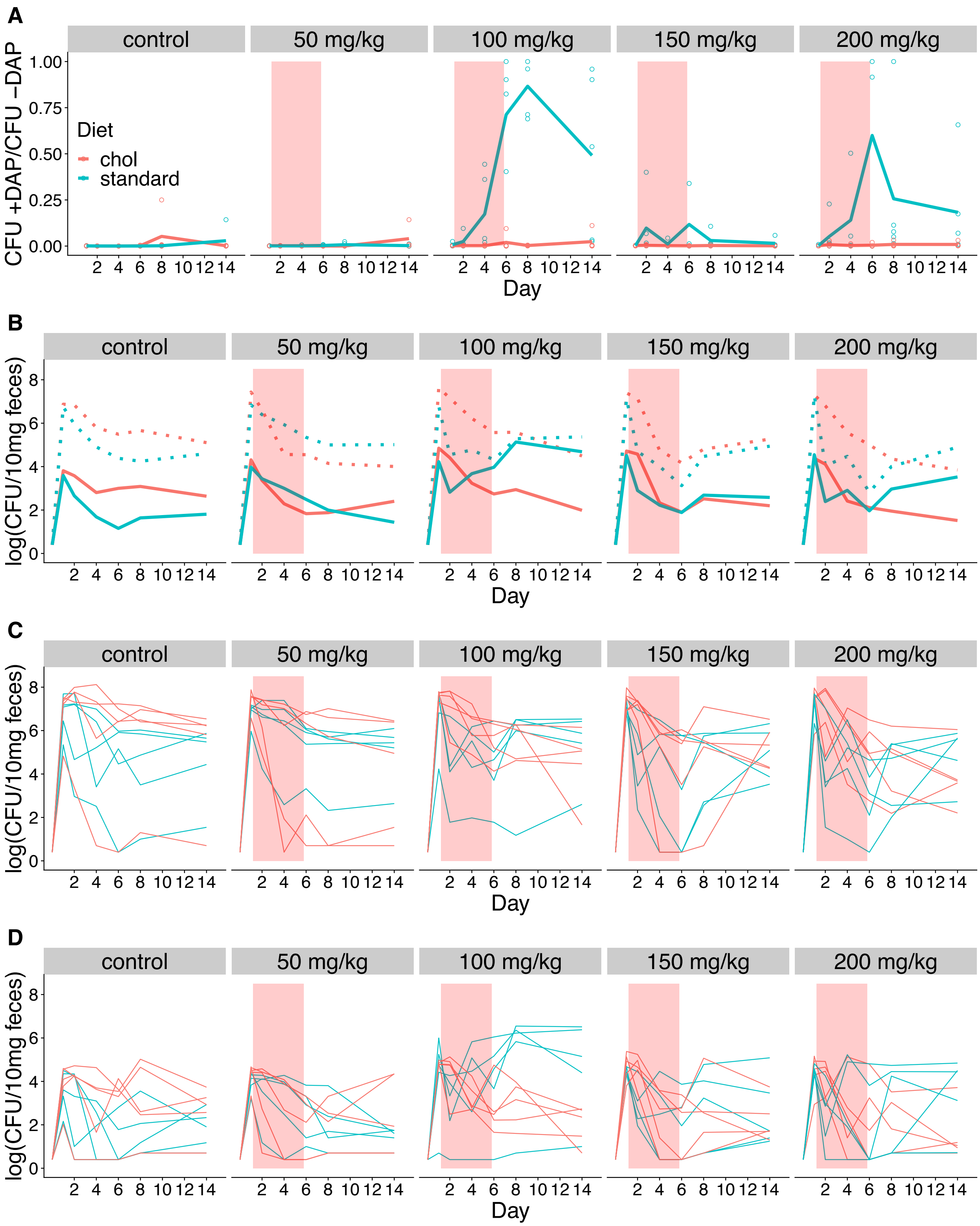


**Figure 5 - figure supplement 3.** Fecal shedding of VR *E.* faecium for Experiment C (strain PR00708-14, Swiss Webster mice, cholestyramine started one day prior to first daptomycin dose). (A) The proportion of fecal VR *E. faecium* that were daptomycin-resistant over time in mice. Proportions were determined by plating on agar with daptomycin (+DAP) and without daptomycin (-DAP). Lines show means, and open points show values for individual mice (N=5). Proportions were not determined for samples with < 20 CFU VR *E. faecium* per 10 mg feces, as these densities were below the limit of detection for this plating assay, and these samples were not included in Panel A. The pink shaded region indicates days of daptomycin therapy. (B) Total VR *E. faecium* densities corresponding to data shown in Panel A (N=5, mean shown). Dotted line shows total density (-DAP) and solid line shows the density of daptomycin-resistant VR *E. faecium* (+DAP). All samples, including those with low densities, were included in Panel B. (C) Total VR *E. faecium* densities (-DAP) for individual mice in this experiment. Each line tracks values for one mouse. (D) Daptomycin-resistant VR *E. faecium* densities (+DAP) for individual mice in this experiment. Each line tracks values for one mouse.

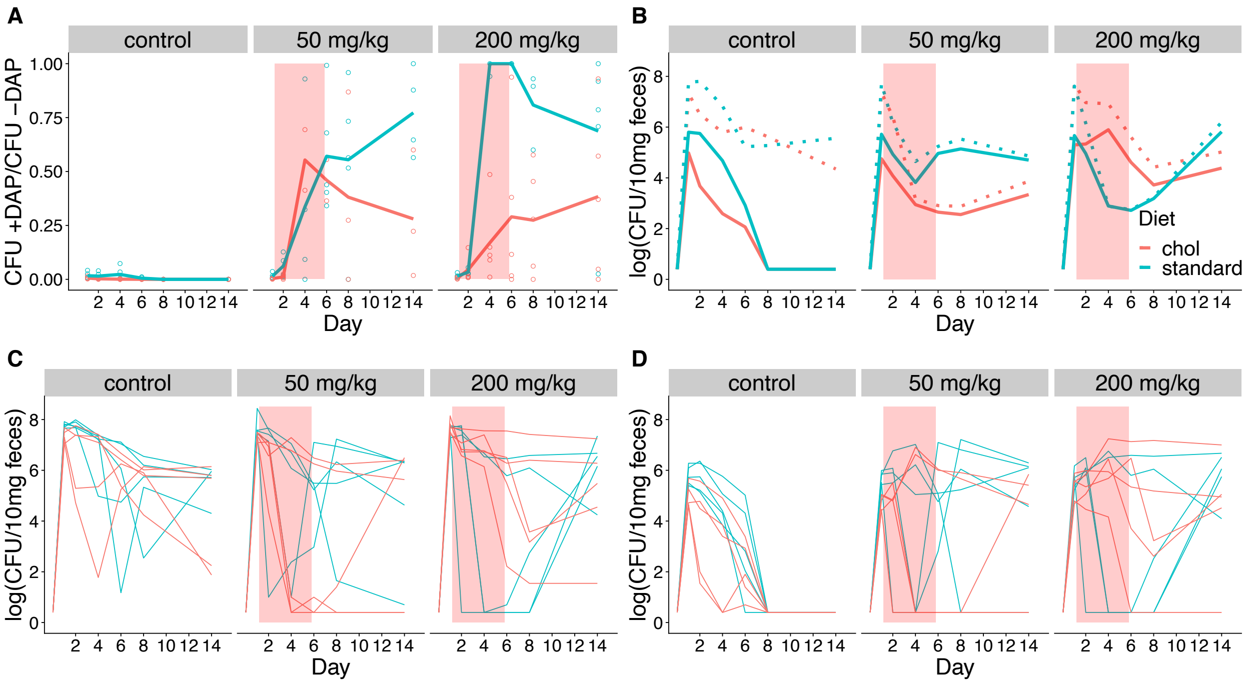


**Figure 5 - figure supplement 4.** Fecal shedding of VR *E.* faecium for Experiment D (strain BL00239-1, Swiss Webster mice, cholestyramine started same day as first daptomycin dose). (A) The proportion of fecal VR *E. faecium* that were daptomycin-resistant over time in mice. Proportions were determined by plating on agar with daptomycin (+DAP) and without daptomycin (-DAP). Lines show means, and open points show values for individual mice (N=5). Proportions were not determined for samples with < 20 CFU VR *E. faecium* per 10 mg feces, as these densities were below the limit of detection for this plating assay, and these samples were not included in Panel A. The pink shaded region indicates days of daptomycin therapy. (B) Total VR *E. faecium* densities corresponding to data shown in Panel A (N=5, mean shown). Dotted line shows total density (-DAP) and solid line shows the density of daptomycin-resistant VR *E. faecium* (+DAP). All samples, including those with low densities, were included in Panel B. (C) Total VR *E. faecium* densities (-DAP) for individual mice in this experiment. Each line tracks values for one mouse. (D) Daptomycin-resistant VR *E. faecium* densities (+DAP) for individual mice in this experiment. Each line tracks values for one mouse.

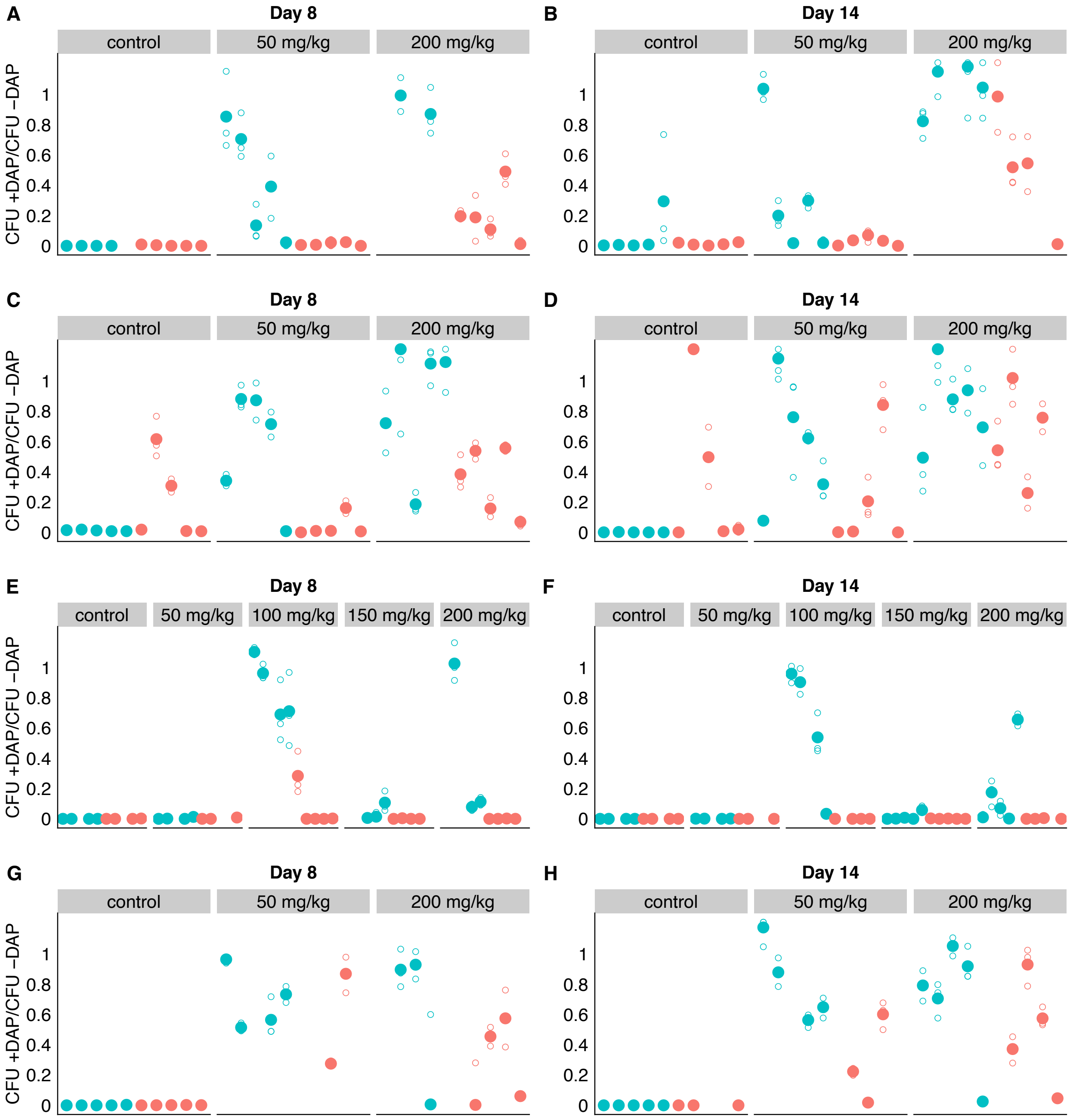


**Figure 5 - figure supplement 5.** Adjunctive cholestyramine prevents emergence of daptomycin resistance in GI tract. Mouse fecal suspensions were plated on Enterococcus-selective plates with daptomycin (+ DAP) and without daptomycin (-DAP) at Day 8 and Day 14 at an estimated 200 CFU per plate (based on previously determined densities). Each filled point represents the mean of triplicate measures from a single mouse sample, and open points show individual measurements. Blue points represent mice fed on a standard diet and red points represent mice fed a cholestyramine-supplemented diet. Mice were treated with daptomycin or saline (controls) for 5 days at the doses listed (N = 5 mice per treatment). Samples with VR *E. faecium* density < 3 x $10^3$ CFU/10 mg feces had insufficient bacterial density to perform this assay, and were not included. Data for three experiments are shown. Values >1 are consistent with sampling variation. **(A-B)** Experiment A. Swiss-Webster mice colonized with strains BL00239-1 + BL00239-1-R, cholestyramine started one day prior to daptomycin. **(C-D)** Experiment B.

C57BL/6 mice colonized with strains BL00239-1 + BL00239-1-R, cholestyramine started one day prior to daptomycin. **(E-F)** Experiment C. Swiss-Webster mice colonized with strains PR00708-14 & PR00708-14-R, cholestyramine started one day prior to daptomycin. **(G-H)** Experiment D. Swiss-Webster mice colonized with strains BL00239-1 + BL00239-1-R, cholestyramine started same day as daptomycin.
